# Uncovering the Hidden Gems of Psoriasis Literature: An Neural Language Model-Assisted Interactive Web Tool

**DOI:** 10.1101/2023.03.15.532867

**Authors:** Sunsi Wu, Xinpei Gu, Ruiheng Xiao, Hongzhi Gao, Bo Yang, Yanlan Kang

## Abstract

**BACKGROUND:** The comprehensive data on psoriasis research are numerous and complex, making it difficult to retrieve and classify manually. The ability to quickly mine literature based on various fine topics using deep learning natural language processing technology to assess research topics and trends in the field of psoriasis disease will have a significant impact on doctors’ research and patients’ health education

**METHOD:** A neural topic model is used to identify fine topics of psoriasis literature published in the PubMed database from 2000 to 2021. Dermatologists evaluate the algorithm-modeled topics, summarize the categories into the most effective topics, and perform linear trend model analysis. The accurate classified topics are presented on an interactive web page to identify research hotspots and trends.

**RESULTS:** At the categorical level, after review by clinicians, 158 out of 160 generated topics were found effective and categorized into 8 groups: Therapeutic methods (34.34%), pathological mechanisms (23.46%), comorbidity (20.04%), Clinical manifestations and differential diagnosis (12.77%), experimental modalities and methods (3.22%), diagnostic tools (2.99%), epidemiology (1.75%), and meetings/guidelines (1.43%). A linear regression model had good accuracy (MSE=0.252602, SSE=42.1845) and strong correlation (R-Squared=0.898009). ANOVA results showed that categories significantly impacted the model (*p*<=0.05), with experimental modalities and methods having the strongest relationship with year, and clinical manifestations and differential diagnosis having the weakest. An interactive web tool (https://psknlr.github.io) facilitates quick retrieval of titles, journals, and abstracts in different categories, as well as browsing literature information under specific topics and accessing corresponding article pages for professional knowledge on psoriasis.

**CONCLUSIONS:** The neural topic model and interactive web tool can effectively identify the research hotspots and trends in psoriasis literature, assisting clinicians and patients in retrieving and comparing pertinent topics and research accomplishments of various years.

## Introduction

Psoriasis is a common, chronic, immune-mediated inflammatory skin disease occurring worldwide and at any age. It is estimated that the prevalence of psoriasis in adults ranges from 0.51% to 11.43%, and in children from 0% to 1.37%(1). Typical clinical manifestations of psoriasis include erythema, scales, pruritus, and involvement of nails. Psoriasis often coexists with other disorders including psoriatic arthritis, cardiometabolic disease, mental disorders, inflammatory bowel disease, malignancies, and infections(2–5). Psoriasis is considered to be associated with high prevalence, chronicity, disfiguration, disability, and comorbidity, which lead to significant negative effects on individuals and society(6–8). Therefore, it is crucial to raise consciousness about the complexity of this disease, the importance of early diagnosis, and the improvement of therapeutic methods.

Numerous articles related to psoriasis exist in PubMed that are hard for dermatologists to read and acquire sufficient information. Most conventional retrieval methods relied on word retrieval and failed to adequately categorize research topics. Medical Subject Headings (MeSH)^(9–11)^ improves retrieval depth by indexing topics with medical terms and a hierarchical tree structure. But it might be hard to accurately classify psoriasis research using the limited medical terminology, which is updated once a year for general diseases. Some secondary words have affected the retrieval system’s evaluation of literature topics, hiding real topic-type data. It is challenging to accurately assess the alterations and trends in research, so it is crucial to focus your search on high-precision topic types.

Natural language processing has become a smooth and efficient way to understand medical domain knowledge, whether in real-time voice or text. The latent Dirichlet allocation (LDA)^(12)^ algorithm is a generative probabilistic clustering model made of a three-level hierarchical Bayesian network. It performs well in text topic classification with a lower perplexity score and higher accuracy. The limitations of LDA are that unsuitable for short-form text^(13)^ which works slowly on noisy and unstructured data^(14)^ and acquire contextual information insufficiently. Though, there are many medical types of research based on LDA^(15–17)^, which phenomenon confirms its universality and efficiency^(18)^ in medical fields. A medical research paper’s abstract is typically a brief text that uses medical jargon and makes strong logical connections, which differs from text in other fields in structure, language, and tone; therefore, LDA may not work on medical topic generation.

As the pre-training language model evolves, inferring the distribution of topics and words becomes more important. How to assign words to biomedical research topics accurately is a new research highlight. In recent years, the transformer-based network inspired by the attention mechanism of brain science and cognitive psychology shows excellent performance in natural language processing and computer vision, such as BERT^(19)^, VIT^(20)^, RoBERTa^(21)^, Sentence-BERT^(22)^, and so on. BERTopic^(23)^, a neural topic model, which leverages Sentence-BERT^(22)^ as word-embedding strategy and class-based TF-IDF for topic representation, has become a popular topic model and will play a vital role in the next few years. The experiment shows that BERTopic gains a dramatic performance^(23)^ among plenty of benchmark models on both short-text corpus and long-text corpus for gaining the highest topic coherence^(24)^ and topic diversity^(25)^.

In this article, we use the BERTopic model to generate topics for 2000-2021 PubMed psoriasis literature. After clinical dermatologists reviewed and summarized the categories, an interactive web tool visualized psoriasis research topics and literature information. so that clinicians can quickly identify topic trends in various categories, search corresponding literature based on topics, and gather enough knowledge from the vast body of literature.

## Methods

### Data Source

We searched PubMed articles between 2000 and 2021 for psoriasis. 38677 relevant articles were retrieved and saved in comma-separated value (CSV) format encoded with UTF-8. After removing duplicate rows and missing rows, 31425 structured psoriasis article entries with title, journal, abstract, publication date, and PubMed ID (PMID) were created.

### Data Preprocessing

All data processing procedures used Python 3.7. The scikit-learn package removed stop words and punctuation. To avoid noise data, we use pandas’ filter and lambda functions to remove special characters like @. Due to the abundance of abbreviations in biomedicine, such as IL-27, we do not handle such cases to ensure the integrity of domain terms in the input data and the interpretability of the output results.

### Neural Topic Model Analyzing

BERTopic is a neural topic model helpful for discovering hot spots and trends in psoriasis topics from a large amount of text. For measuring semantic similarity between weighted topics, BERTopic mainly uses Sentence-BERT^(22)^ word embedding, UMAP^(26)^ dimensionality reduction, DBSCAN^(27)^ clustering, and class-based TF-IDF to model the topic:

Assuming there is a document set *D*, each document *d_i_* ∈ *D* consists of *n_i_* sentences 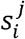, where *j* ∈ [1, *n_i_*], each sentence 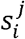 consists of 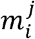 words 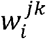, where 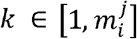. First, use the BERT-Sentence model to map each sentence 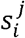 to a word embedding space 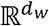, where *d_w_* is the word embedding dimension, that is, 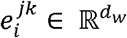, where 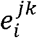 is the word embedding vector of 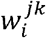. Then, sum the word embedding vectors 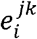 of each sentence 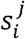 get the sentence embedding vector 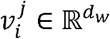, that is:

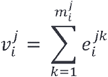

Then, use UMAP to reduce the sentence embedding vector 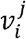 to 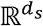, where *d_s_* is the sentence embedding dimension, that is, 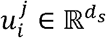, where 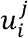 is the sentence embedding vector of 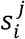. Then, use the DBSCAN algorithm to cluster the sentence embedding vector 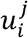, and get *K* clusters *C_k_*, where *k* ∈ [1, *K*]. Finally, use the class-based TF-IDF algorithm to model each cluster *C_k_*, and get the topic model *M_k_*, where *M_k_* represents the topic model of *C_k_*, that is:

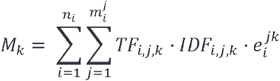

where *TF_i,j,k_* is the term frequency of 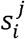 in the cluster *C_k_, IDF_i,j,k_* is the inverse document frequency of 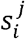 in the cluster *C_k_*, and 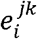 is the word embedding vector of 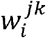.

### Classification of Topics

Dermatologists usually have stronger classification abilities in few-shot learning than algorithmic models. For further analysis of the trend of topic distribution, quantities of the representation of topics were manually divided into several categories. Unimportant, hard-to-classify topics must be removed.

### Linear Trend Model

The trend line model is a linear regression analysis of the relationship between the publication year and the logarithmic count of psoriasis papers, taking into account the categorical factor. The category significance is tested at a 0.05 level of significance.

## Results

### Topic and Classification Results

160 topics were extracted and recognized by the neural topic model and 158 (98.75%) effective psoriasis topics (eTable1) were identified after removing the −1st and 112th topics. These topics were displayed on an intertopic distance map (Fig 2) and represented using word clouds (eFigure1), and were divided into 8 categories (eTable1) by dermatologists for analysis of psoriasis research trends and highlights: comorbidities, therapeutic methods, pathological mechanisms, diagnostic tools, clinical manifestations and differential diagnosis, meetings and guidelines, experimental modalities and methods, and epidemiology.

**Fig 1.**
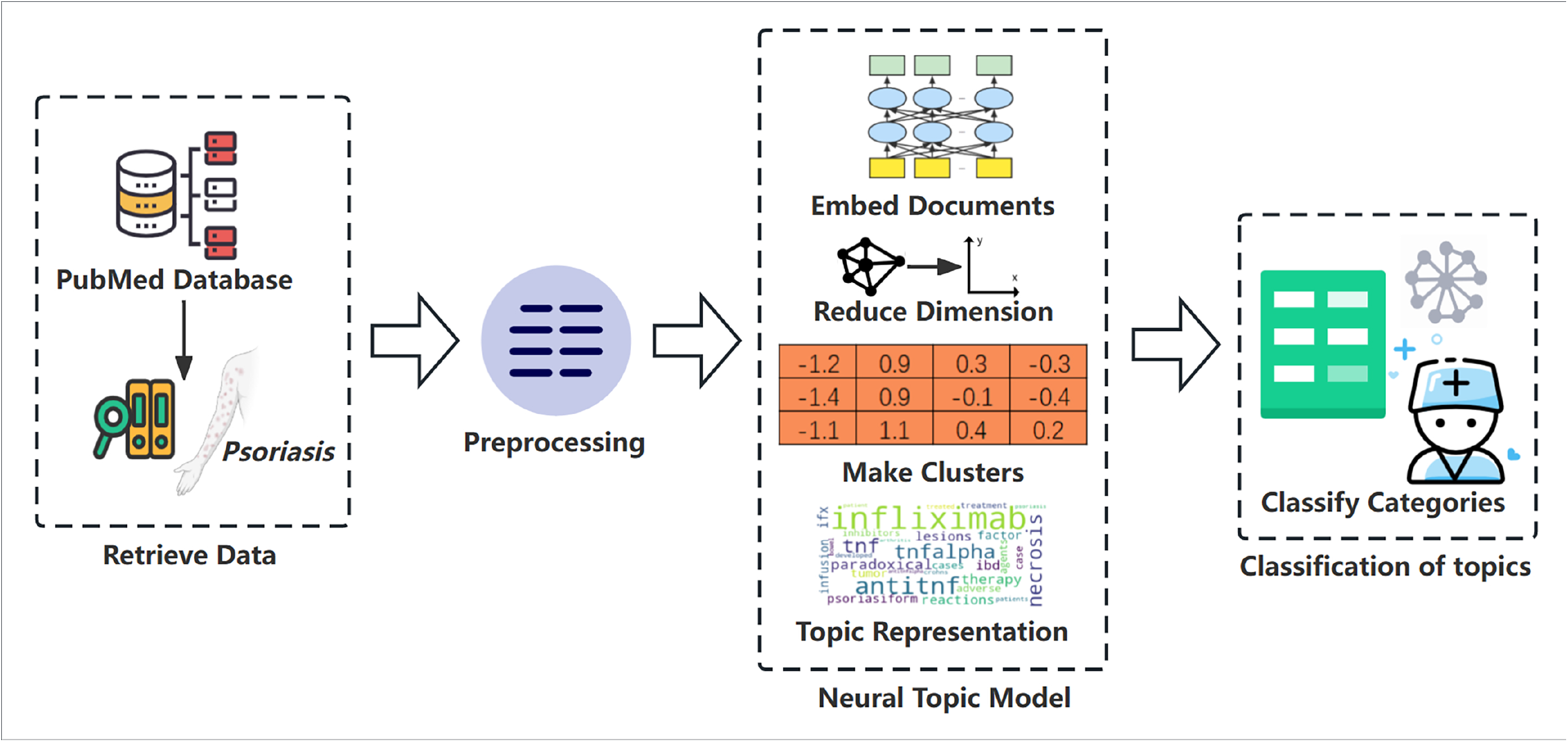
Flow chart for analyzing psoriasis research hotspots and trends. After gathering and preprocessing publication data, generating document embedding, and then making clusters and producing topic representation based on a neural topic model.

**Fig 2.**
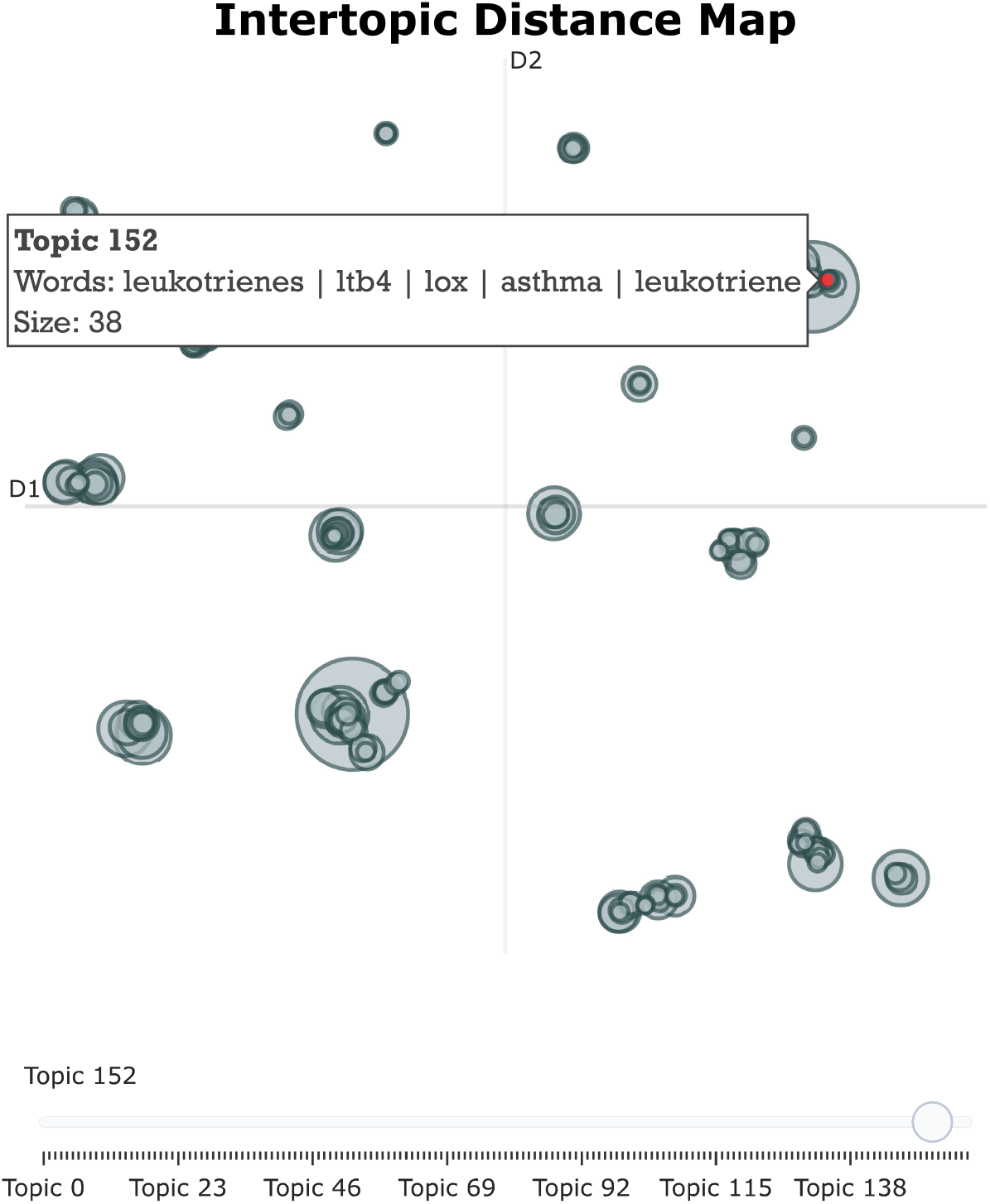
Intertopic distance map of 163 psoriasis topics in 2D representation. The interactive graph can reveal the core words of the topic and its size

**Fig 3.**
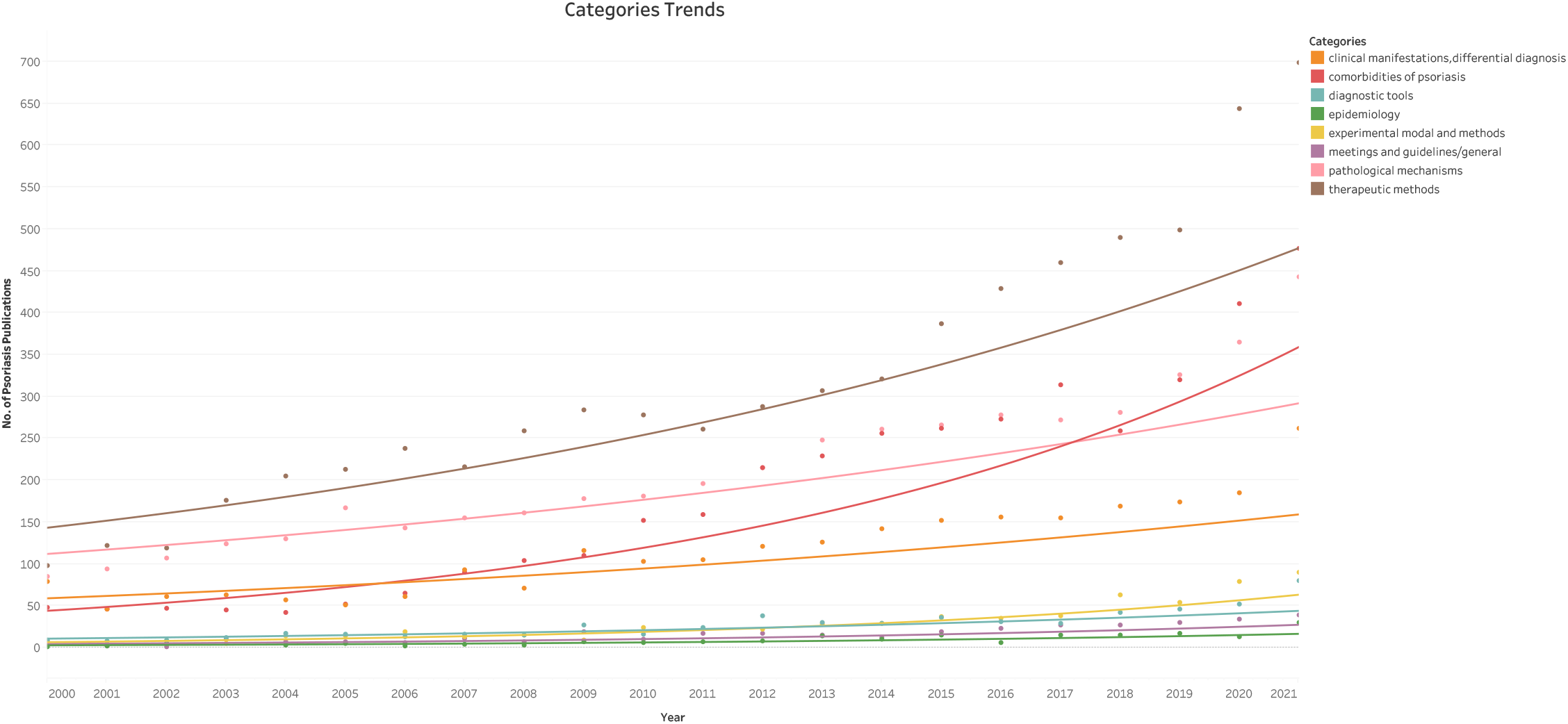
Category trends based on linear regression model. The scatter plot depicts the correlation between the number of articles published in various years and their corresponding categories, utilizing a linear regression model to extrapolate the underlying trend.

**Fig 4.**
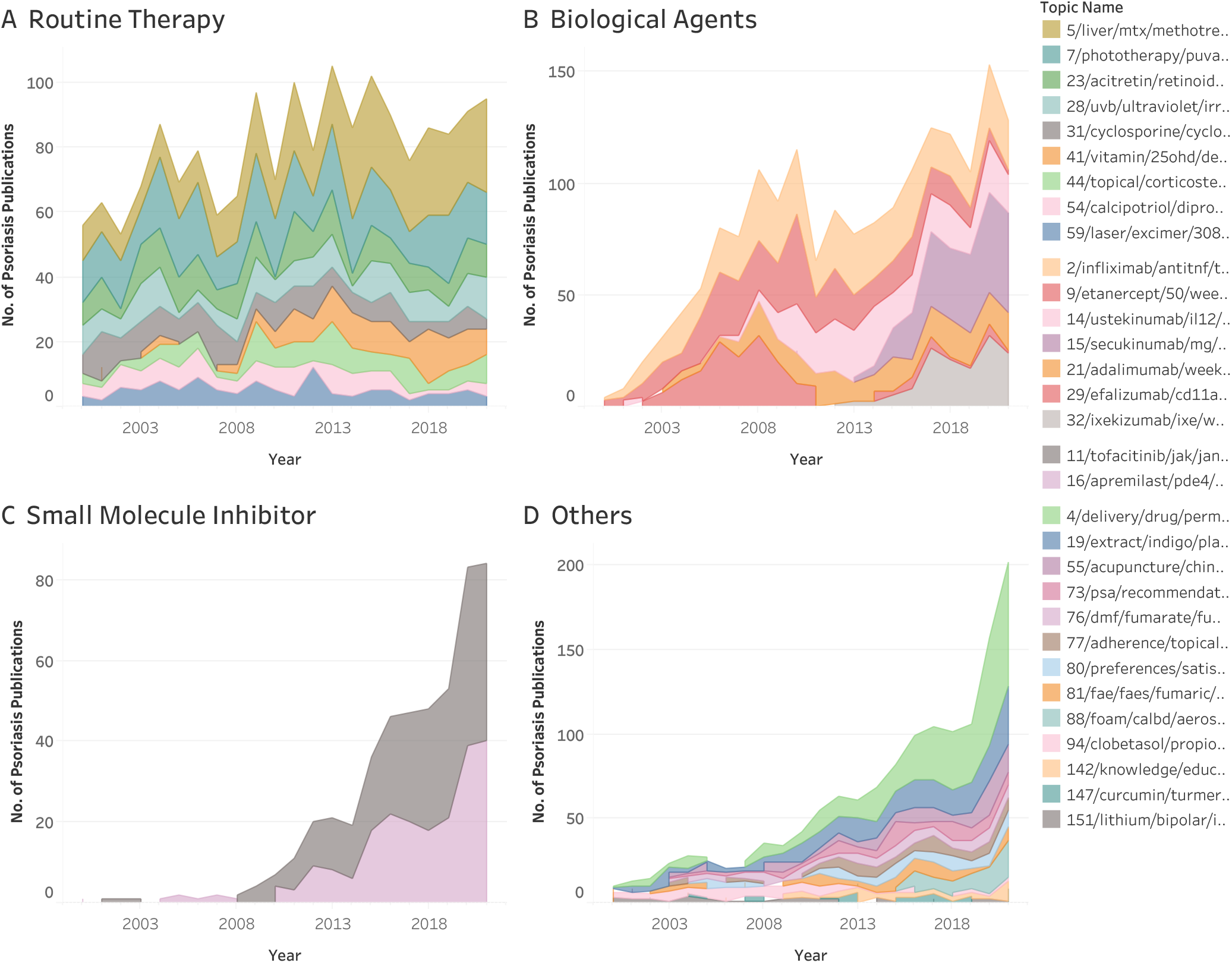
Topic trends in therapeutic methods on interactive psoriasis literature website. The information including the specific topic, paper title, abstract, publication date, and PubMed ID (PMID) can be viewed on the interactive web tool (https://psknlr.github.io)

### Linear Trend Model Results

The linear regression model examines the relationship between psoriasis paper counts and year, with 183 observations. The formula is “Categories * (log of year + intercept)” and has a high prediction accuracy (MSE=0.252602, SSE=42.1845). The R-Squared value of 0.898009 indicates a strong correlation and the ANOVA shows that categories significantly impact the model (*p*<=0.05). The “experimental modalities and methods” category has the strongest relationship with year, while “clinical manifestations, differential diagnosis” has the weakest.

### Interactive Websites of Neural Topic Model

We created a visual and interactive website based on Tableau to ensure convenience and traceability of psoriasis paper data. There are basic functions and advanced functions for web tool. The basic operation level allows quick switching and viewing of topics across classifications and searching for literature by title, journal, and abstract. The advanced function can precisely locate the title, release date, journal, and category details of the literature corresponding to a given year and related topics. The detailed operation procedure is illustrated in the appendix. The distribution and trend of some important categories will be described below.

## Research Topics and Trends

### Field of Clinical Manifestations and Differential Diagnosis

The clinical psoriasis field was divided into three subcategories: clinical classification, clinical manifestations, and differential diagnoses. In the clinical classification subcategory, nail psoriasis (17.03%) was the most researched topic, followed by pediatric psoriasis (7.72%) and pregnancy psoriasis (7.14%). Generalized pustular psoriasis (2.87%) emerged as a research topic since 2011. In the clinical manifestations subcategory, pruritus (4.54%), Koebner (2.33%), and geographic tongue (2.17%) were the top three research topics. In differential diagnoses, spondyloarthritis (SpA) (13.34%) and atopic dermatitis (AD) (5.86%) were the most researched topics, reaching their peak in 2021. Inflammatory linear verrucous epidermal nevus (ILVEN) (5.35%) and acute generalized exanthematous pustulosis (AGEP) (3.92%) had a stable number of publications.

### Field of Pathological Mechanisms and Comorbidities

Three subcategories were classified based on pathological mechanisms. The majority of topics were divided into subcategories of inheritance and immunity and inflammation, with the remainder falling under “others.” There were 3 notable trends. Firstly, in the inheritance subcategory, genetics-related publications (22.83%) were at a high level and peaked in 2015. The trend of microRNA variation (6.38%) has risen significantly since 2016. Recent studies on epigenetic methylation (1.67%) and mesenchymal stem cells (0.91%) have also been conducted. Secondly, despite slight fluctuations, the majority of topics in the immunity and inflammation subcategory showed consistent trends, with gut microbiota (7.86%) showing consistent growth since 2017 and studies on T help 17 cells/interleukin-17 axis (2.47%) and interleukin-36 (1.08%) becoming stable after 2012. Lastly, in other subcategories, the number of research papers on most topics showed slight fluctuations. In the field of psoriasis comorbidity, most topics showed an upward trend after 2013, with psoriasis and cardiovascular disease (40.34%) leading and psoriatic arthritis (6.50%) ranking second. The number of pandemic disease (4.62%) publications, which fluctuated between 0 and 6 before, increased tenfold between 2020 and 2021, causing a noticeable shift in the field.

### Field of Therapeutic Methods

We divided it into four subcategories, with a focus on routine therapy and biological therapies. Routine therapy showed a stable number of publications, with methotrexate (5.60%) as the leading academic paper since 2014. Phototherapy (5.18%) ranked second, with a higher number of studies prior to 2010. Biological therapies showed fluctuations: infliximab (6.33%), etanercept (4.72%), and ustekinumab (3.75%) showed similar variations with peaks in 2008, 2010, and 2012, respectively. Efalizumab (2.64%) showed an upward trend before 2006 but was suspended from the market in 2009 due to complications(28, 29). Secukinumab (3.40%) and ixekizumab (2.08%) grew rapidly to become the most popular monoclonal antibodies since 2017. In the small molecule inhibitors subcategory, tofacitinib (4.05%) and apremilast (3.25%) showed noticeable growth. Lipid-based nanoparticles (5.60%) were the most researched route of administration since 2010, and calcipotriol/betamethasone dipropionate aerosol foam was a fresh and efficient technique since 2015.

## Discussion

The conventional topic modeling techniques can be divided into two groups: bag-of-words models, which extract topics based on word co-occurrence frequency but ignore contextual semantics. The other is clustering based on a pre-trained language model to make word embedding, which can lead to centroid-based inductive bias(25, 30). BERTopic model brings new inspiration to solve the problem of incompatibility between clustering and extraction of words, and formed a special neural topic modeling way, making the extraction of core words suitable for irregularly shaped clusters in medical scenes. Of note, we also found that sometimes the key factor related to psoriasis can be extracted, such as HLAB27-induced axial spondyloarthritis, which means the neural topic model can learn the knowledge representation of psoriasis to a certain extent.

Conventional topic modeling techniques can be categorized into two groups: bag-of-words models and clustering-based models. The former extract topics based solely on word co-occurrence frequency, ignoring contextual semantics. The latter utilize pre-trained language models to generate word embeddings, which can lead to a centroid-based inductive bias. The BERTopic model presents a novel solution to the challenge of reconciling clustering and word extraction by employing a specialized neural topic modeling approach. This allows for the extraction of core words in irregularly shaped clusters in medical settings. Our findings also indicate that the neural topic model can learn a representation of knowledge related to psoriasis to some degree, as demonstrated by its ability to extract key factors, such as HLAB27-induced axial spondyloarthritis.

Currently available tools, such as PubMed, rely heavily on word retrieval in the abstract. Nonetheless, the central idea of literature cannot be captured by a single term. The incorporation of secondary words into some literature abstracts influences the retrieval system’s evaluation of literature topics, obscuring the discovery of the true topic of the literature. Our study employs a neural topic model to mine the literature for deep semantic information. This method models the topic using the total weight of all abstract vocabulary vectors and TF-IDF, which lets us find relevant literature quickly and accurately. The current retrieval system encourages reinforcing former knowledge. Although the artificial formulation of multiple retrieval strategies based on Boolean logic operators(31) can improve understanding of research accomplishments within the current disease knowledge framework, this can come at the expense of missing out on mining some particularly rich disease-related subject topics and emerging rare topics. We use dermatologists to filter and classify the topics generated by the algorithm, which is displayed with a visual web tool to ensure the ease of use and professionalism of disease literature retrieval. Clinicians can benefit from knowing the historical context of disease research and the most recent clinical advancements. Likewise, normal readers can quickly build a logical framework and acquire information about diseases without the need for sophisticated retrieval methods or specialized knowledge.

Our study revealed that psoriasis had changed dramatically over the past 21 years in terms of pathogenesis, diagnosis, and treatment. The in-depth study of the mechanism clarifies psoriasis classification and management, especially for children and pregnant women. Pustular forms are separated from plaque psoriasis as molecular and genomic insights add to clinical phenotype(32). The research topic of generalized pustular psoriasis (GPP) had gotten more attention since 2011 when IL-36RN mutations playing a pivotal role in the pathogenesis of GPP were found. The issuance of relevant guidelines has made pediatric psoriasis a hot topic in the last five years. The National Psoriasis Foundation (NPF) released the screening guidelines for children with psoriasis complications in 2017(33). The American Academy of Dermatology (AAD) in conjunction with the NPF published the nursing management and treatment of pediatric psoriasis in 2019(34). The research topic of pregnancy had become popular since 2018 with a sharp growth. These newly added publications were mainly concerned with biological agents in the evaluation and management of pregnant women with psoriasis. The treatment paradigm for psoriasis has indeed changed dramatically in the last 20 years due to the advent of biologics. There are currently 11 biologics approved by the US Food and Drug Administration (FDA) for the treatment of psoriasis with good efficacy and few adverse effects. However, even with the rapid development of biologics, conventional systemic therapy is still recommended as the first line of treatment for psoriasis by various guidelines. Conventional systemic therapy remains unique in its affordability, oral-only administration, and known safety profile. New drugs such as PDE4 inhibitors, JAK inhibitors, TYK2 inhibitors, and others can be taken orally or applied topically without injections, giving psoriasis patients more options(35–37). Clinical trials of more small molecule drugs are currently underway, and in the future, there is an urgent need to obtain high-quality clinical data to better guide clinical practice. The optimization of dosage forms, such as nanoformulations, guarantees the safety of the drug, reduces toxic side effects, and improves clinical efficacy. Meanwhile, the inclusion of psoriasis comorbidity management further refines the management paradigm.

In 2010, World Health Organization (WHO) statistics indicate that psoriasis possesses the highest mortality rates despite diabetes and hypertension having the highest prevalence of coronary artery disease and vascular lesions. Cardiovascular disease was the leading cause of death among psoriasis patients(38). Patients with psoriatic arthritis may experience swelling, pain, deformation, and even disability in their joints, which can seriously affect their daily life(39). To take both skin symptoms and comorbid diseases into consideration, the treatment framework is shifting from short-term intervention for acute rashes to long-term management and early diagnosis and treatment can help improve the quality of life of patients. More than that, psoriasis and many comorbidities raise the risk of mental illnesses such as anxiety and depression in psoriasis patients(40). These emotional problems will aggravate skin lesions and form a vicious circle. Education is key for these patients. It provides psychological relief, shifts the psoriasis treatment paradigm to a new model of physician-patient collaboration, and promotes patient understanding of clinical treatment strategies. Overall, the complexities of psoriasis are slowly being unraveled, and our neural topic model and retrieval method provide the convenience of simplifying the complex for both clinicians and normal readers.

## Limitation

Though neural topic models have reshaped the topic modeling method and show extraordinary performance, which can process documents much faster than humans, their understanding ability through analytic tasks such as classification still needs to be improved, and the comprehension to domain knowledge is still inferior to that of medical experts. It is still necessary to analyze the professional literature utilizing neural network rapid processing and manual interpretation by medical experts. The topic representation may consist of a small amount of redundant information caused by the neural topic model.

## Conclusion

The neural topic model enables the extraction of core words appropriate for irregularly shaped clustering clusters in medical scenes. Key topic words related to psoriasis publications can be extracted, which means the pre-trained language model can learn the knowledge representation of psoriasis to a certain extent. Mining the semantic data of the psoriasis research literature based on topic depth, in conjunction with the review and classification of clinicians, can quickly and professionally understand the relevant research under the fine classification of psoriasis clinical manifestations and differential diagnosis, pathological mechanisms, and treatment methods. Likewise, create a visual article retrieval system to make it simpler and more intuitive for medical professionals and patients to understand how research is progressing.

**Table 1.** Linear trend model analysis of various categories in psoriasis research. Analysis of the impact of year on various categories of psoriasis research, including therapeutic methods, pathological mechanisms, meetings and guidelines, experimental models and methods, epidemiology, diagnostic tools, comorbidities, and clinical manifestations. Results show significant linear trends with p-values less than 0.05.

**No funding sources**

## Conflicts of Interest

The authors declare that the research was conducted in the absence of any commercial or financial relationships that could be construed as a potential conflict of interest.

## Data Availability Statement

The research’s original contributions are listed in the article/Supplemental Material; further questions can be directed to the corresponding author.

## Author Contributions

Dr. Y. Kang and Xiao had full access to all of the data in the study and take responsibility for the integrity of the data and the accuracy.

Concept and design: SW, YK and BY.

Acquisition, analysis, or interpretation of data: SW, RX and YK.

Drafting of the manuscript: SW and XG.

Statistical analysis: YK, XG, RX, HG.

Obtained funding: None.

Administrative, technical, or material support: YK, RX, HG.

Supervision: SW, YK, BY, RX, HG and XG.

The published version of the manuscript has been read and approved by all authors.

## Supplementary Material

The Supplementary Material for this article can be found online at:

(https://psknlr.github.io)

## SUPPLEMENT 1

1. eMethods: Stopwords removed.
2. eFigure1: Word clouds.
3. eTable1: List of topics from the topic model.
4. eFigure2: Web Page Usage.
5. eFigure3: Tendency chart.

## Notes

### Competing Interest Statement

The authors have declared no competing interest.

