## Supplementary material for "Uncovering the Hidden Gems of Psoriasis Literature: An Neural Language Model-Assisted Interactive Web Tool": Table 1

Table 1. Linear trend model analysis of various categories in psoriasis research

| Categories | p-value | DF | Term | Value | StdErr | t-value | p-value |
| --- | --- | --- | --- | --- | --- | --- | --- |
| therapeutic methods | 0.0001808 | 21 | Year | 0.0001573 | 3.47E-05 | 4.53514 | 0.0001808 |
|  |  |  | intercept | -0.786596 | 1.40903 | -0.558252 | 0.582576 |
| pathological mechanisms | 0.0015129 | 21 | Year | 0.0001254 | 3.44E-05 | 3.6451 | 0.0015129 |
|  |  |  | intercept | 0.13173 | 1.39736 | 0.0942701 | 0.925788 |
| meetings and guidelines/general | 0.0010265 | 21 | Year | 0.000254 | 6.67E-05 | 3.8083 | 0.0010265 |
|  |  |  | intercept | -7.94342 | 2.70925 | -2.93196 | 0.0079657 |
| experimental modal and methods | < 0.0001 | 21 | Year | 0.0003072 | 3.54E-05 | 8.66824 | < 0.0001 |
|  |  |  | intercept | -9.43994 | 1.43934 | -6.55851 | < 0.0001 |
| epidemiology | < 0.0001 | 20 | Year | 0.0002594 | 5.13E-05 | 5.05736 | < 0.0001 |
|  |  |  | intercept | -8.70255 | 2.09025 | -4.1634 | 0.0004801 |
| diagnostic tools | < 0.0001 | 21 | Year | 0.0001897 | 3.78E-05 | 5.02037 | < 0.0001 |
|  |  |  | intercept | -4.61352 | 1.53436 | -3.0068 | 0.0067167 |
| comorbidities of psoriasis | < 0.0001 | 21 | Year | 0.0002753 | 3.97E-05 | 6.93508 | < 0.0001 |
|  |  |  | intercept | -6.28496 | 1.61223 | -3.89829 | 0.0008283 |
| clinical manifestations,differential diagnosis | 0.0024999 | 21 | Year | 0.0001304 | 3.80E-05 | 3.43247 | 0.0024999 |
|  |  |  | intercept | -0.697543 | 1.54306 | -0.452052 | 0.655866 |
