## SUPPLEMENT 1 for "Uncovering the Hidden Gems of Psoriasis Literature: An Neural Language Model-Assisted Interactive Web Tool"

**APPENDIX – SUPPLEMENTARY MATERIALS**

Table of Contents

1. Stopwords removed

2. Word clouds

3. List of topics from the topic model.

4. Web Page Usage

5. Tendency chart

### 1. Stopwords removed

**sklearn package:**

“a”

“about”

“across”

“afterwards”

“again”

“all”

“almost”

“alone”

“along”

“although”

“also”

“already”

“always”

“am”

“among”

“amongst”

“amount”

“amoungst”

“an”

“and”

“any”

“anything”

“anyway”

“anywhere”

“are”

“as”

“at”

“back”

“be”

“became”

“because”

“become”

“becomes”

“becoming”

“been”

“before”

“beforehand”

“behind”

“being”

“below”

“between”

“beside”

“besides”

“beyond”

“bill”

“both”

“bottom”

“but”

“by”

“call”

“can”

“cannot”

‘cant’

“co”

“con”

“could”

“couldn’t”

“cry”

“de”

“detail”

“describe”

“do”

“done”

“down”

“due”

“during”

“each”

“eg”

“eight”

“either”

“eleven”

“else”

“elsewhere”

“empty”

“enough”

“even”

“ever”

“every”

“everything”

“etc”

“except”

“few”

“fifteen”

“fifty”

“fill”

“find”
“fire”
“first”
“five”
“for”
“former”
“formerly”
“forty”
“found”
“four”
“from”
“front”
“full”
“further”
“get”
“give”
“go”
“had”
“has”
“hasn’t”
“have”
“he”
“hence”
“her”
“here”
“herein”
“hereupon”
“hers”
“herself”
“him”
“himself”
“his”
“how”
“however”
“hundred”
“i”
“ie”
“if”
“in”
“inc”
“indeed”
“interest”
“is”
“it”

“its”

“itself”

“keep”

“last”

“latter”

“latterly”

“least”

“less”

“ltd”

“made”

“may”

“me”

“meanwhile”

“might”

“mill”

“mine”

“my”

“myself”

“more”

“move”

“most”

“much”

“must”

“name”

“namely”

“neither”

“never”

“nevertheless”

“next”

“nobody”

“none”

‘noone’

“not”

“nor”

“nothing”

“nowhere”

“of”

“off”

“often”

“on”

“once”

“one”

“only”

“or”

“other”
“others”
“otherwise”
“our”
“ours”
“ourselves”
“out”
“over”
“own”
“part”
“per”
“perhaps”
“please”
“put”
“rather”
“re”
“same”
“see”
“seem”
“seemed”
“seeming”
“seems”
“serious”
“several”
“she”
“should”
“show”
“side”
“since”
“sincere”
“six”
“sixty”
“so”
“some”
“somehow”
“someone”
“sometime”
“sometime”
“sometimes”
“somewhere”
“still”
“such”
“system”
“take”

“ten”
“than”
“that”
“the”
“their”
“them”
“themselves”
“then”
“thence”
“there”
“thereafter”
“thereby”
“therefore”
“therein”
“thereupon”
“these”
“they”
“thick”
“thin”
“third”
“this”
“those”
“though”
“three”
“through”
“throughout”
“thru”
“thus”
“to”
“together”
“too”
“top”
“toward”
“towards”
“twelve”
“twenty”
“two”
“un”
“under”
“until”
“up”
“upon”
“us”
“very”

“via”
“was”
“we”
“well”
“were”
“what”
“whatever”
“when”
“whence”
“whenever”
“where”
“whereafter”
“whereas”
“whereby”
“wherein”
“whereupon”
“wherever”
“whether”
“which”
“while”
“whither”
“who”
“whoever”
“whole”
“whom”
“whose”
“why”
“will”
“with”
“within”
“without”
“would”
“yet”
“you”
“your”
“yours”
“yourself”
“yourselves”

**nltk package:**
“a”
“about”
“again”
“ain”

“all”

“am”

“an”

“and”

“any”

“are”

‘aren’

“aren’t”

“as”

“be”

“because”

“been”

“before”

“being”

“below”

“between”

“both”

“but”

“by”

“can”

“could”

“couldn’t”

“d”

‘doesn’

“doesn’t”

“did”

‘didn’

“didn’t”

“do”

“does”

“doing”

“down”

“during”

“each”

“few”

“for”

“from”

“further”

“had”

“hadn”

“hadn’t”

“have”

‘haven’

“haven’t”

“having”
“has”
‘hasn’
“hasn’t”
“he”
“her”
“here”
“hers”
“herself”
“him”
“himself”
“his”
“how”
“i”
“if”
“in”
“into”
“it”
“it’s”
“its”
“is”
‘isn’
“isn’t”
“just”
“l”
“m”
“ma”
“me”
‘mightn’
“mightn’t”
“more”
“most”
‘mustn’
“mustn’t”
“my”
“myself”
‘needn’
“needn’t”
“no”
“nor”
“now”
“not”
“o”
“of”

“off”
“on”
“once”
“only”
“or”
“other”
“our”
“ours”
“ourselves”
“out”
“over”
“own”
“re”
“s”
“same”
“shan”
“shan‘t”
“she”
“she‘s”
“should”
‘shouldn‘
“shouldn‘t”
“should‘ve”
“so”
“some”
“such”
“t”
“than”
“that”
“that‘ll”
“the”
“their”
“theirs”
“them”
“themselves”
“then”
“there”
“these”
“they”
“this”
“those”
“through”
“to”
“too”

“under”

“untill”

“up”

“ve”

“very”

“was”

‘wasn’

“wasn‘t”

“we”

“were”

‘weren’

“weren’t”

“what”

“when”

“where”

“which”

“while”

“who”

“whom”

“why”

“will”

“with”

“won”

“won’t”

‘wouldn’

“wouldn’t”

“y”

“you”

“you’d”

“you’ll”

“your”

“you’re”

“yours”

“yourselves”

“you’ve”

### 2. Word clouds

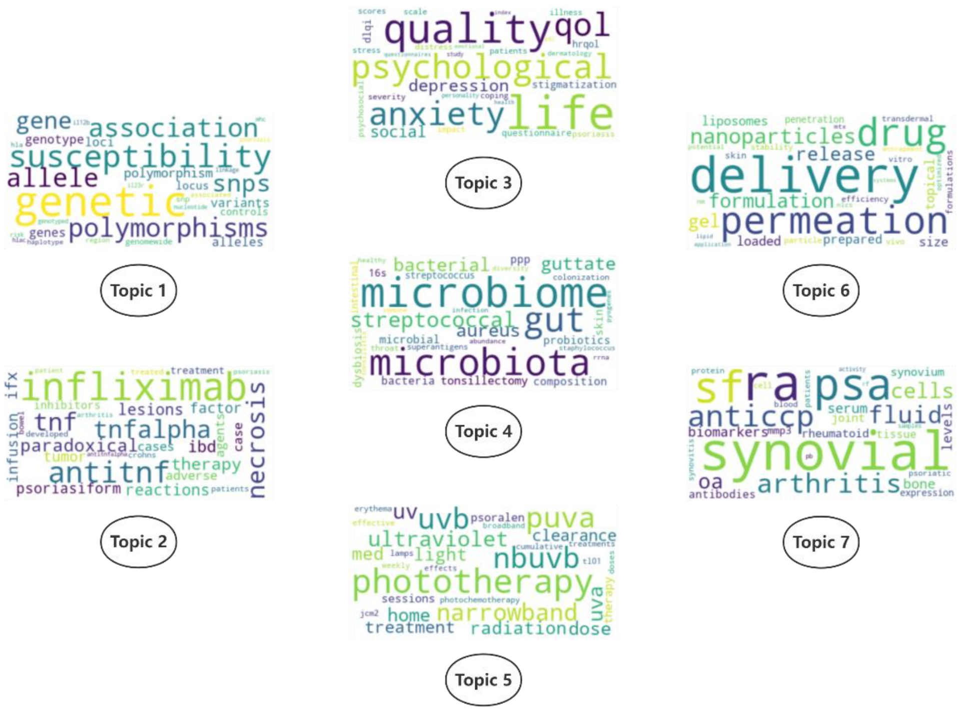


Word clouds for top 7 topics of psoriasis publication published from 2000 to 2021. The higher the weight of the core words, the larger the size of the word possesses.

### 3. List of topics from the topic model, and associated categorisations.

| Aggregated topic | Subcategory | ID | Topic name | Count | Proportion |
|---|---|---|---|---|---|
| Comorbidities of psoriasis | | 0 | cardiovascular/risk/metabolic/obesity | 1632 | 40.37% |
| | | 12 | activity/psa/mda/measures | 263 | 6.51% |
| | | 24 | tb/tuberculosis/ltbi/tst | 192 | 4.75% |
| | | 26 | covid19/pandemic/vaccination/sarscov2 | 187 | 4.63% |
| | | 36 | hepatitis/hcv/hbv/infection | 143 | 3.54% |
| | | 42 | stigmatization/psychological/social/anxiety | 131 | 3.24% |
| | | 46 | autoimmune/thyroid/ms/thyroiditis | 121 | 2.99% |
| | | 89 | screening/pase/pest/psa | 72 | 1.78% |
| | | 95 | sleep/osa/rls/psqi | 68 | 1.68% |
| | | 103 | renal/ckd/nephropathy/kidney | 66 | 1.63% |
| | | 51 | synovial/ra/sf/psa | 120 | 2.97% |
| | | 57 | ocular/uveitis/eye/eyes | 107 | 2.65% |
| | | 71 | mf/fungoides/mycosis/lymphoma | 90 | 2.23% |
| | | 75 | hiv/infection/antiretroviral/aids | 84 | 2.08% |
| | | 78 | alopecia/aa/hair/areata | 85 | 2.10% |
| | | 84 | bullous/pemphigoid/bp/pemphigus | 77 | 1.90% |
| | | 87 | vitiligo/repigmentation/segmental/mktp | 74 | 1.83% |
| | | 110 | ibd/uc/cd/eims | 58 | 1.43% |
| | | 113 | bone/bmd/osteoporosis/density | 58 | 1.43% |
| | | 117 | hpv/hhv7/dna/hpv5 | 54 | 1.34% |
| | | 139 | cancer/melanoma/nmsc/risk | 42 | 1.04% |
| | | 145 | hz/zoster/herpes/vaccine | 41 | 1.01% |
| | | 150 | psychiatric/disorders/psychocutaneous/trichotillomania | 38 | 0.94% |
| | | 152 | leukotrienes/ltb4/lox/asthma | 37 | 0.92% |
| | | 131 | periodontitis/periodontal/salivary/probing | 42 | 1.04% |
| | | 123 | melanoma/cd147/pkm2/traf6 | 49 | 1.21% |
| | | 121 | sarcoidosis/pneumonia/granulomatous/granulomas | 52 | 1.29% |
| | | 109 | sexual/ed/genital/erectile | 60 | 1.48% |
| Therapeutic methods | Biological Agents | 2 | infliximab/antitnf/tnfalpha/paradoxical | 439 | 6.33% |
| | Others | 4 | delivery/drug/permeation/nanoparticles | 388 | 5.60% |
| | Routine Therapy | 5 | liver/mtx/methotrexate/nafld | 388 | 5.60% |
| | Routine Therapy | 7 | phototherapy/puva/nbuvb/uvb | 359 | 5.18% |
| | Biological Agents | 9 | etanercept/50/weekly/mg | 327 | 4.72% |
| | Small Molecule Inhibitor | 11 | tofacitinib/jak/janus/inhibitors | 281 | 4.05% |
| | Biological Agents | 14 | ustekinumab/il12/mg/45 | 260 | 3.75% |
| | Biological Agents | 15 | secukinumab/mg/300/150 | 236 | 3.40% |
| | Small Molecule Inhibitor | 16 | apremilast/pde4/phosphodiesterase/inhibitor | 225 | 3.25% |
| | Others | 19 | extract/indigo/plant/plants | 215 | 3.10% |
| | Biological Agents | 21 | adalimumab/week/ada/safety | 198 | 2.86% |
| | Routine Therapy | 23 | acitretin/retinoids/retinoid/isotretinoin | 199 | 2.87% |
| | Routine Therapy | 28 | uvb/ultraviolet/irradiation/uv | 180 | 2.60% |
| | Biological Agents | 29 | efalizumab/cd11a/itolizumab/humanized | 183 | 2.64% |
| Therapeutic methods | Routine Therapy | 31 | cyclosporine/cyclosporin/csa/cya | 159 | 2.29% |
| | Biological Agents | 32 | ixekizumab/ixe/weeks/week | 144 | 2.08% |
| | Biological Agents | 37 | brodalumab/il17/210/mg | 144 | 2.08% |
| | Biological Agents | 39 | tnfi/persistence/psa/retention | 135 | 1.95% |
| | Routine Therapy | 41 | vitamin/25ohd/deficiency/d3 | 133 | 1.92% |
| | Routine Therapy | 44 | topical/corticosteroids/corticosteroid/steroids | 127 | 1.83% |
| | Biological Agents | 45 | alefacept/ach/courses/course | 126 | 1.82% |
| | Routine Therapy | 54 | calcipotriol/dipropionate/betamethasone/twocompound | 115 | 1.66% |
| | Others | 55 | acupuncture/chinese/medicine/chm | 113 | 1.63% |
| | Routine Therapy | 59 | laser/excimer/308nm/pdl | 110 | 1.59% |
| | Biological Agents | 61 | survival/drug/biologic/ustekinumab | 94 | 1.36% |
| | Biological Agents | 64 | guselkumab/week/voyage/100 | 94 | 1.36% |
| | Routine Therapy | 72 | tacrolimus/pimecrolimus/topical/ointment | 89 | 1.28% |
| | Others | 73 | psa/recommendations/arthritis/treattotarget | 88 | 1.27% |
| | Routine Therapy | 74 | pdt/photodynamic/ala/ppix | 85 | 1.23% |
| | Others | 76 | dmf/fumarate/fumaric/mmf | 86 | 1.24% |
| | Others | 77 | adherence/topical/medication/nonadherence | 77 | 1.11% |
| | Others | 80 | preferences/satisfaction/attributes/preference | 78 | 1.13% |
| | Others | 81 | fae/faes/fumaric/esters | 77 | 1.11% |
| | Biological Agents | 82 | biosimilars/biosimilar/reference/originator | 79 | 1.14% |
| | Others | 88 | foam/calbd/aerosol/dipropionate | 75 | 1.08% |
| | Others | 94 | clobetasol/propionate/foam/spray | 74 | 1.07% |
| | Routine Therapy | 98 | vitamin/vdr/d3/analogs | 70 | 1.01% |
| | Biological Agents | 99 | iraes/checkpoint/nivolumab/immunotherapy | 70 | 1.01% |
| Therapeutic methods | Routine Therapy | 105 | tazarotene/hptaz/lotion/hp | 63 | 0.91% |
| | Biological Agents | 108 | agents/efalizumab/alefacept/biologic | 61 | 0.88% |
| | Routine Therapy | 111 | cyclosporine/therapies/methotrexate/agents | 59 | 0.85% |
| | Routine Therapy | 115 | leflunomide/mtx/lef/methotrexate | 54 | 0.78% |
| | Biological Agents | 126 | risankizumab/90/150/16 | 47 | 0.68% |
| | Routine Therapy | 128 | ointment/calcitriol/tacalcitol/microgg | 49 | 0.71% |
| | Biological Agents | 133 | tildrakizumab/resurface/200/100 | 43 | 0.62% |
| | Biological Agents | 136 | czp/certolizumab/400/pegol | 45 | 0.65% |
| | Others | 142 | knowledge/educational/health/care | 41 | 0.59% |
| | Others | 147 | curcumin/turmeric/curcuma/spice | 37 | 0.53% |
| | Others | 151 | lithium/bipolar/inositol/blockers | 37 | 0.53% |
| | Others | 153 | psa/arthritis/agents/drugs | 37 | 0.53% |
| | Routine Therapy | 154 | sea/dead/salt/uvb | 38 | 0.55% |

| | | | | | |
|---|---|---|---|---|---|
| Pathological mechanisms | Immunity and Inflammation | 56 | keratinocytes/il17a/k17/expression | 115 | 2.43% |
| | Others | 58 | oxidative/antioxidant/stress/levels | 108 | 2.28% |
| | Others | 63 | stress/depression/cortisol/hpa | 94 | 1.99% |
| | Immunity and Inflammation | 66 | chemokine/cxcr3/chemokines/ccr4 | 98 | 2.07% |
| | Others | 67 | antimicrobial/amps/cathelicidin/peptides | 96 | 2.03% |
| | Others | 68 | platelet/mpv/ykl40/crp | 89 | 1.88% |
| | Others | 69 | alcohol/smoking/consumption/smokers | 95 | 2.01% |
| | Inheritance | 79 | methylation/epigenetic/dna/modifications | 79 | 1.67% |
| | Immunity and Inflammation | 83 | dcs/lcs/dendritic/dc | 79 | 1.67% |
| | Immunity and Inflammation | 92 | il20/il22/il19/il24 | 75 | 1.58% |
| | Immunity and Inflammation | 93 | angiogenesis/antiangiogenic/growth/vegf | 71 | 1.50% |
| | Immunity and Inflammation | 101 | mast/crh/cells/cell | 66 | 1.39% |
| | Immunity and Inflammation | 102 | psoriasin/s100a7/s100a8/s100 | 66 | 1.39% |
| | Others | 119 | ppar/pioglitazone/peroxisome/ppars | 54 | 1.14% |
| | Immunity and Inflammation | 120 | pdcs/dendritic/plasmacytoid/Il37 | 53 | 1.12% |
| | Immunity and Inflammation | 122 | il36/il36r/il36ra/cytokines | 51 | 1.08% |
| | Immunity and Inflammation | 125 | cd/celiac/iga/antibodies | 51 | 1.08% |
| | Others | 127 | ros/oxidative/redox/ho1 | 49 | 1.03% |
| | Others | 130 | zinc/zn/copper/iron | 48 | 1.01% |
| | Others | 132 | adhesion/leukocyte/integrins/selectins | 45 | 0.95% |
| | Immunity and Inflammation | 134 | serum/cytokines/levels/ifn | 46 | 0.97% |
| | Inheritance | 135 | mscs/mesenchymal/stem/msc | 43 | 0.91% |
| | Immunity and Inflammation | 137 | rort/ror/inverse/orphan | 42 | 0.89% |
| | Inheritance | 143 | bcl2/apoptosis/apoptotic/p53 | 42 | 0.89% |
| Pathological mechanisms | Others | 148 | metabolites/metabolomic/metabolomics/metabolite | 38 | 0.80% |
| | Others | 149 | n3/fatty/pufa/acids | 38 | 0.80% |
| | Others | 155 | sensory/nerve/neuropeptides/cgrp | 36 | 0.76% |
| | Immunity and Inflammation | 157 | spa/il23/il17/il17a | 36 | 0.76% |
| Diagnostic tools | | 13 | images/rcm/skin/imaging | 265 | 43.87% |
| | | 43 | enthesitis/ultrasound/entheses/entheseal | 129 | 21.36% |
| | | 49 | mri/imaging/bone/synovitis | 120 | 19.87% |
| | | 114 | machine/prediction/gene/network | 54 | 8.94% |
| | | 158 | dermoscopic/dermoscopy/pr/vessels | 36 | 5.96% |
| Clinical manifestations, differential diagnosis | Clinical Classification | 3 | nail/onychomycosis/nails/napsi | 439 | 17.03% |
| | Differential Diagnosis | 8 | spa/axial/spondyloarthritis/hlab27 | 344 | 13.34% |
| | Clinical Classification | 22 | children/pediatric/childhood/adolescents | 199 | 7.72% |
| | Clinical Classification | 27 | pregnancy/women/birth/pregnancies | 184 | 7.14% |
| | Differential Diagnosis | 33 | ad/dermatitis/dermatology/dermatological | 151 | 5.86% |
| | Clinical Classification | 34 | ppp/palmoplantar/pustulosis/soles | 147 | 5.70% |
| | Differential Diagnosis | 40 | linear/verrucous/ilven/lesions | 138 | 5.35% |
| | Clinical Manifestation | 50 | pruritus/itch/intensity/symptom | 117 | 4.54% |
| | Differential Diagnosis | 62 | agep/pustular/generalized/pustules | 101 | 3.92% |
| | Differential Diagnosis | 85 | contact/dermatitis/patch/allergens | 78 | 3.03% |
| | Clinical Classification | 86 | gpp/il36rn/pustular/mutation | 74 | 2.87% |
| | Clinical Classification | 90 | involvement/arthritis/joint/joints | 76 | 2.95% |
| | Differential Diagnosis | 96 | synovial/sf/ra/cells | 70 | 2.72% |
| | Differential Diagnosis | 97 | lichen/genital/sclerosus/vulvar | 70 | 2.72% |
| | Clinical Manifestation | 107 | koebner/tattoo/phenomenon/tattoos | 60 | 2.33% |
| Clinical manifestations, differential diagnosis | Clinical Manifestation | 116 | tongue/oral/gt/geographic | 56 | 2.17% |
| | Clinical Classification | 118 | jia/juvenile/ilar/jpsa | 55 | 2.13% |
| | Differential Diagnosis | 124 | tinea/trichophyton/capitis/incognito | 53 | 2.06% |
| | Differential Diagnosis | 129 | sapho/osteomyelitis/crmo/osteitis | 48 | 1.86% |
| | Differential Diagnosis | 140 | malassezia/species/yeasts/globosa | 41 | 1.59% |
| | Clinical Classification | 144 | erythroderma/erythrodermic/etiology/diagnosis | 42 | 1.63% |
| | Differential Diagnosis | 156 | anticcp/antibodies/rf/ra | 35 | 1.36% |
| Meetings and guidelines/general | | 18 | guidelines/systemic/consensus/dermatologists | 225 | 78.13% |
| | | 106 | grappa/meeting/research/annual | 63 | 21.88% |
| Experimental modal and methods | | 20 | 20/mice/imqinduced/imiquimod/psoriasislike | 214 | 32.97% |
| | | 30 | 30/life/quality/dlqi/qol | 172 | 26.50% |
| | | 60 | 60/pasi/dlqi/pga/bsa | 105 | 16.18% |
| | | 91 | 91/nma/trials/nmas/network | 74 | 11.40% |
| | | 138 | 138/models/model/animal/mouse | 45 | 6.93% |
| | | 146 | 146/children/cdlqi/life/quality | 39 | 6.01% |
| Epidemiology | | 65 | 65/cost/costs/costeffectiveness/costeffective | 96 | 27.12% |
| | | 100 | 100/costs/cost/economic/health | 67 | 18.93% |
| | | 70 | 70/prevalence/age/incidence/psoriasis | 89 | 25.14% |
| | | 104 | 104/mortality/hazard/95/risk | 60 | 16.95% |
| | | 141 | 141/psa/incidence/prevalence/population | 42 | 11.86% |

### 4. Web Page Usage

In the «http://psknlr.github.io/» web page, switch to the subdivision theme (for example: D. Diagnostic Tools. C. Meetings and Guidelines. And so on.), each small box represents each specific document, and different colors represent different themes. Moving the mouse to each small box can present the information, and clicking it can jump to the PubMed page where the document is located in the new TAB page.

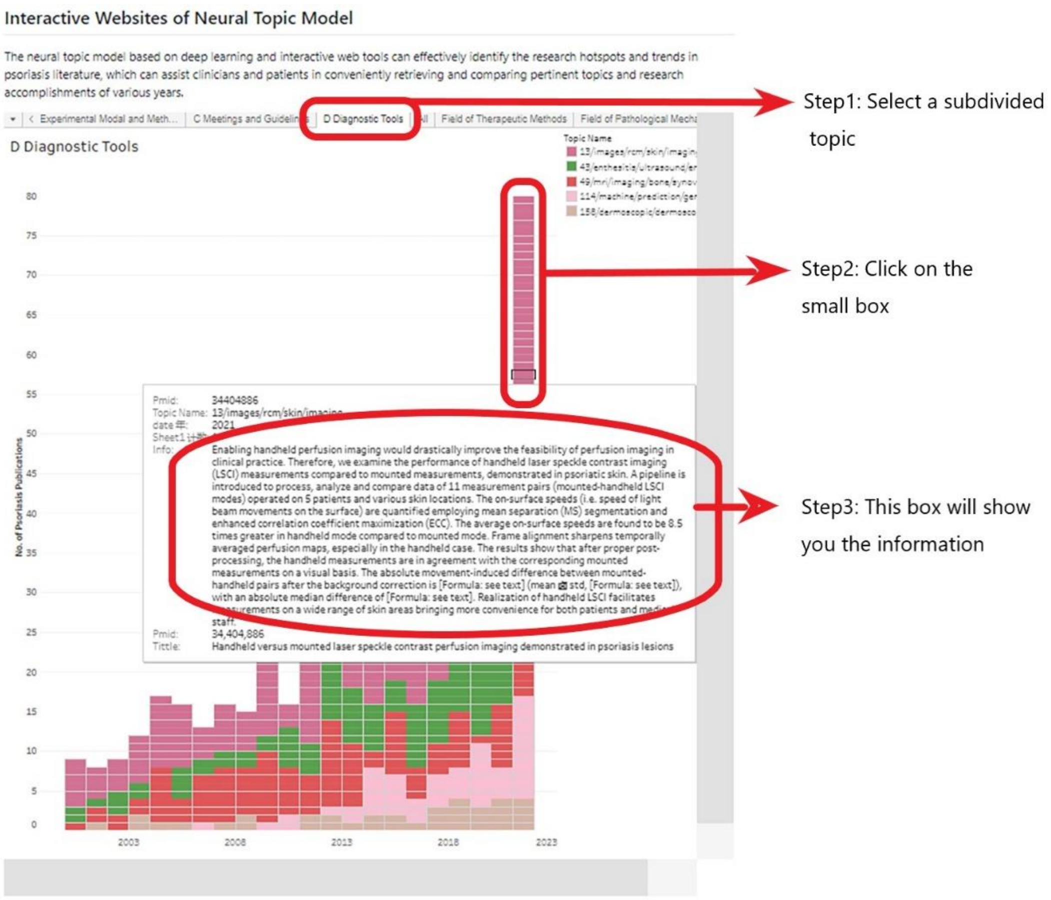

### 5. Tendency chart

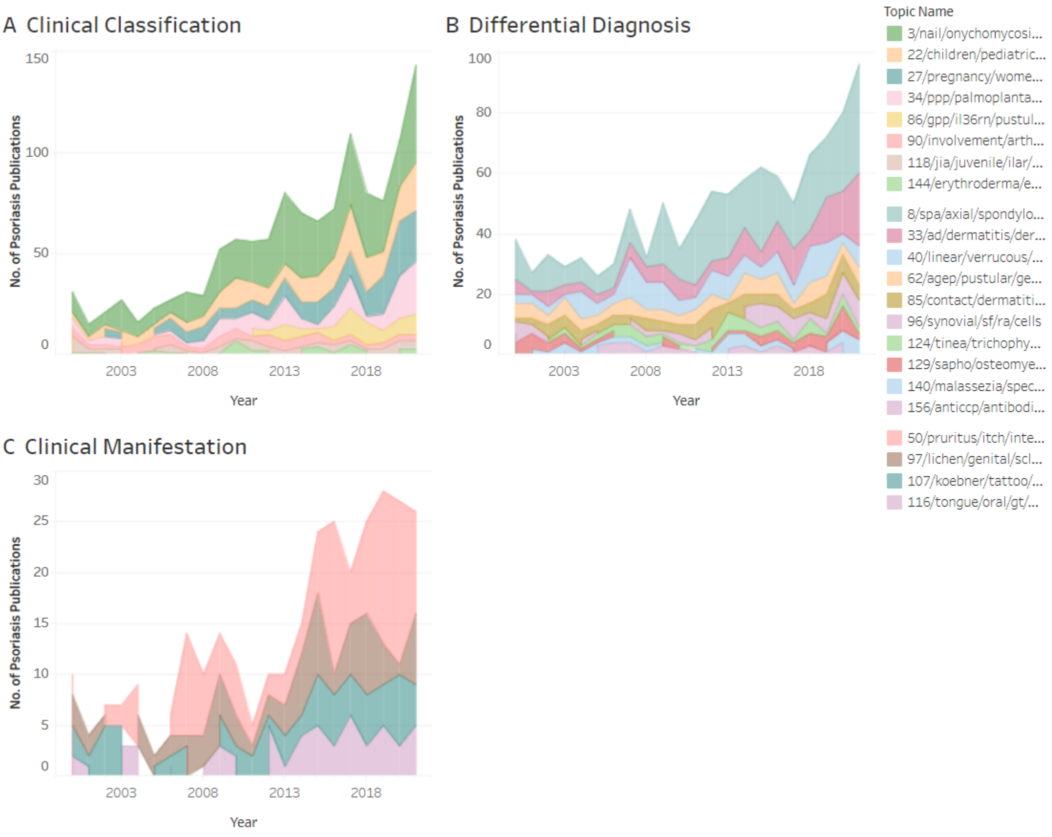


Topic trends in clinical manifestations and differential diagnosis of psoriasis papers published

from 2000 to 2021.

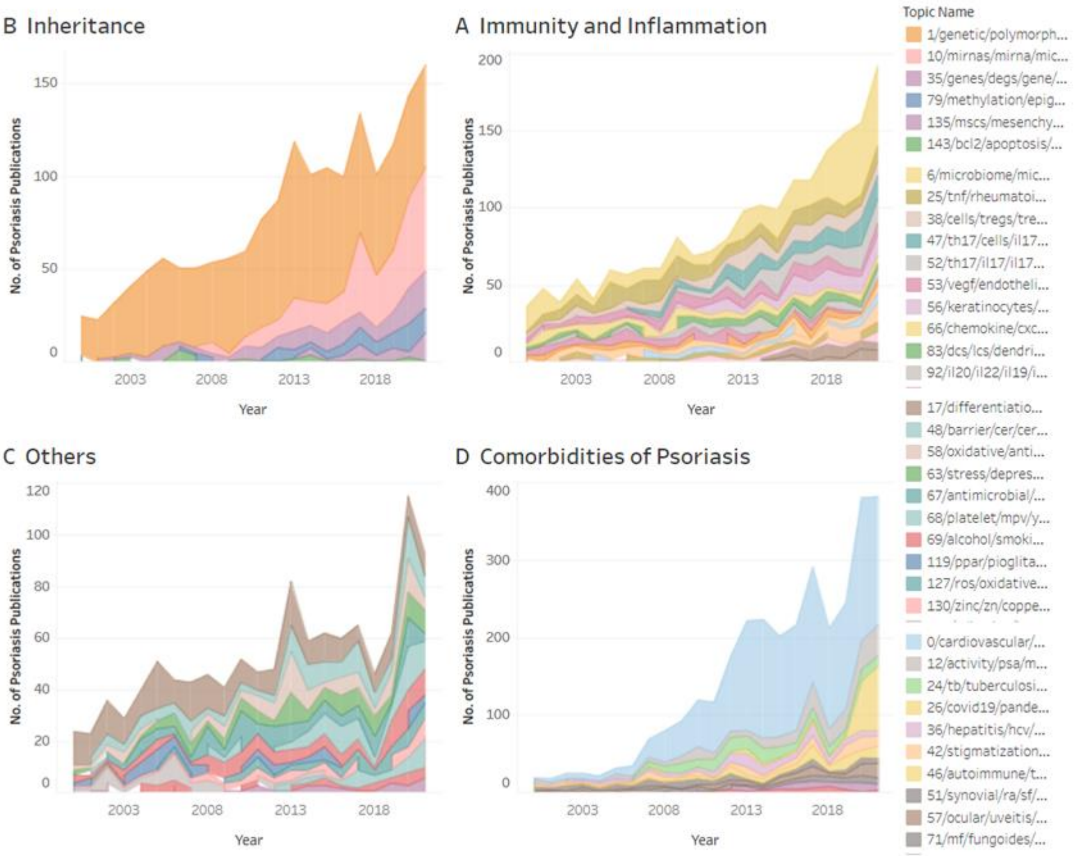


Topic trends in pathological mechanisms and comorbidities of psoriasis papers published from 2000 to 2021.

Each subcategory and comprehensive category can be displayed on the web page, and researchers can click the tab bar above to switch to the specific classification view. At the level of fundamental operations, researchers can click a topic at a specific year on the graph to view the details of the corresponding articles for that topic. The prompt box will then show the classification, topic name, year, and number of articles. When performing advanced operations, selecting a specific era and associated theme, waiting a short while, then clicking the prompt box's top right corner to reveal the associated logo will reveal the complete data pop-up window. The data summary comes first, and it is consistent with the data that the fundamental functions have provided. By the time you click the "complete data" button, the details of the particular article, including the title, publish date, journal, categories, and topic name, will show up.
